# Real-time spike sorting platform for high-density extracellular probes with ground-truth validation and drift correction

**DOI:** 10.1101/101030

**Authors:** James J. Jun, Catalin Mitelut, Chongxi Lai, Sergey L. Gratiy, Costas A. Anastassiou, Timothy D. Harris

## Abstract

Electrical recordings from a large array of electrodes give us access to neural population activity with single-cell, single-spike resolution. These recordings contain extracellular spikes which must be correctly detected and assigned to individual neurons. Despite numerous spike-sorting techniques developed in the past, a lack of high-quality ground-truth datasets hinders the validation of spike-sorting approaches. Furthermore, existing approaches requiring manual corrections are not scalable for hours of recordings exceeding 100 channels. To address these issues, we built a comprehensive spike-sorting pipeline that performs reliably under noise and probe drift by incorporating covariance-based features and unsupervised clustering based on fast density-peak finding. We validated performance of our workflow using multiple ground-truth datasets that recently became available. Our software scales linearly and processes up to 1000-channel recording in real-time using a single workstation. Accurate, real-time spike sorting from large recording arrays will enable more precise control of closed-loop feedback experiments and brain-computer interfaces.

## Introduction

Advancements in our understanding of neural computation and how it leads to complex behavior critically depends on accurate measurements of coordinated neural activities in behaving animals. This technical challenge requires a combination of sensor hardware and analysis software that are well-integrated and optimized for specific signals of interest. Parallel developments in sensor hardware and analysis software are necessary to realize the full potential of the measurement system to meet quality standards.

Extracellular electrodes such as silicon probes and tetrodes offer many recording sites that capture action potentials (spikes) from proximal neurons outside of their cell membranes. Typically, each electrode detects spikes from multiple neurons, which, in a subsequent step, must be correctly assigned to individual neurons. This is possible due to unique spatiotemporal electrical patterns generated by these same neurons as they have different morphology, ion-channel distributions, and spatial positions relative to the electrodes [1]. These factors critically contribute to features of the extracellular spike waveforms at each recording site, which provide useful characteristics for discriminating spikes from different neurons [Henze et al, 2000; Gold et al, 2006; Anastassiou et al, 2015]. Spike-sorting analysis describes the task of assigning individual neuronal identities to each detected spike, and the importance of this computation has increased due to the wide accessibility of extracellular recording technology and the increased requirements to record and separate activity from as many neurons as possible.

Advances in semiconductor fabrication technology permitted developments of high-definition (HD) silicon probes offering a very large number (~1000) of densely packed sites (Figure 1A, A’) [2–6]. These improvements have dramatically increased the recording volume from brain tissue and the spatial resolution of electrical measurements. Furthermore, miniaturized HD silicon probes, which integrate on-chip electronics with recording sites, are quickly becoming commercially available. For example, IMEC Neuropixels probes provide up to 966 densely-packed sites (20 μm vertical site spacing) on a single shank (9.6 mm length) [4,5]. Such probes have enabled monitoring from the entire rodent brain depth during free behaviors, while inflicting minimal tissue damage attributed to a shank occupying a minimal cross-sectional area (70×20 μm^2^).

**Figure 1.**
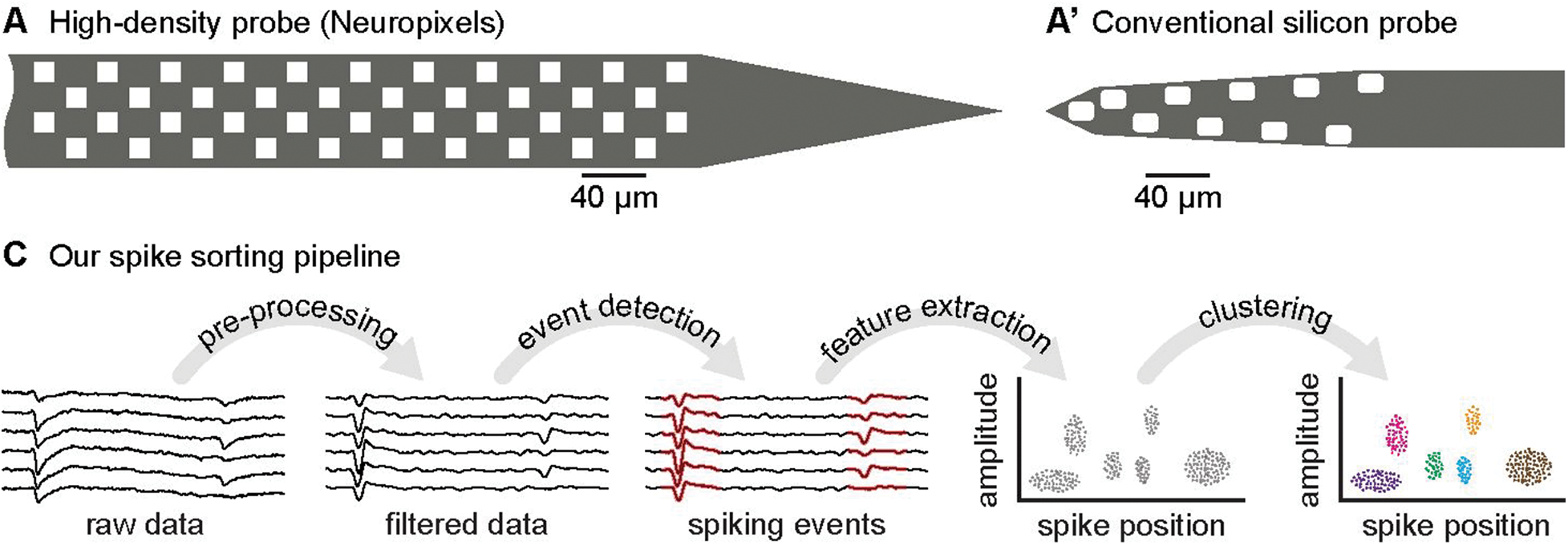
Overview of our spike sorting pipeline. (**A**) High-density silicon probes such as the Neuropixels probes are becoming widely available, which features 960 sites along 9.6 mm shank length (70 μm width). (**A’**) Existing spike-sorting methods are designed for conventional silicon probes having a lower site density and limited length coverage. (**B**) Our proposed spike-sorting pipeline for high-density probes employs algorithms optimized for high-density site patterns with an extended length coverage. One of our key features is an extensive use of the spatial information in spike waveforms from closely-spaced sites. Our sorting pipeline incorporates the spike positions to improve the spike sorting accuracy and to correct for the probe drifts.

These HD probes bring both unique opportunities and challenges for spike sorting analysis. The increased site density can improve the spike sorting accuracy through enhanced spatial information, since there are more sites within the spike detection range. Prevailing spike sorting algorithms [7,8] do not computationally scale well in light of the increasing number of sites on the HD probes and alternative approaches must be considered to accurately and efficiently sort spikes from these probes.

Realistic experimental conditions such as probe spatial drift and noise contamination result in assignment errors that have, up to now, required time-consuming manual corrections. This manual burden in the analysis pipeline has greatly limited the utility of HD probes. Additionally, it is common to observe probe drift due to brain movements, which can cause a spiking neuron (unit) to move out of the probe’s detection range. HD probes can potentially solve this problem by offering extended spatial coverage to track drifting units. By taking advantage of recording hardware as such, it became possible to achieve fully-automated and unsupervised spike sorting to the level of human accuracy or greater.

In this paper, we introduce a comprehensive spike sorting pipeline (*JRCLUST*: Janelia Rocket Clust) that improves on existing techniques by addressing accuracy, speed and usability by exploiting recent advancements in clustering and GPU computing. Our analysis pipeline involves a series of automated stages including pre-processing, event detection, feature extraction and clustering (Figure 1B), followed by an efficient manual verification stage guided by a graphical user interface. Our runtime scales linearly to the data volume, and achieves real-time performance for 1000 channels by efficiently distributing the spike data to parallel processors according to their spatial positions. The global probe movement is automatically detected and corrected using the spatial correlation of spike positions across time. We evaluated the accuracy of JRCLUST using three types of ground-truth datasets specific to HD probes obtained from manual curation [9], paired juxtacellular recordings [10] and biophysical simulations [11].

## Results

### Pre-processing and noise-cancellation strategies during experimental recordings

Extracellular recordings are typically contaminated by various types of noise in experimental settings, and the quality of spike sorting critically depends on the ability to selectively remove the noise without affecting the signal. We consider noise as unwanted signals that corrupt the spike waveforms of interest, and noise is injected by a coupling mechanism unique to their origins and characteristics. We introduce noise cancellation methods to address three of the most common noise types observed during *in-vivo* recordings. In particular, we address the noise sources intrinsic to the brain such as overlapping and background spikes, and external noise sources such as electrical interferences and motion artifacts.

Spike waveforms can be isolated from the raw recordings by passing a frequency range (300 - 6000 Hz) containing high signal powers relative to the noise and LFP powers. However, the resulting waveforms are prone to overlapping spikes due to coincident firing of multiple units in close proximity. It is possible to reduce the occurrence of overlapping spikes by narrowing the width of the spike waveforms using a slope filter (Figure 2A,A’). The slope filter results in narrower spike waveforms (~50% FWHM) since it amplifies the brief initial phase of the action potential dominated by sodium channel dynamics. We apply Savitzky-Golay filter (2^nd^ order, 9 taps) to perform smoothing and differentiation operations in one pass, which is much faster than typical band-pass filter operations requiring an extra data format conversion and two passes in opposite time directions (see Methods). Due to the built-in smoothing operation, our slope filter implementation does not significantly change the SNR of spike waveforms compared to the bandpass filters (Figure 2A’’).

**Figure 2.**
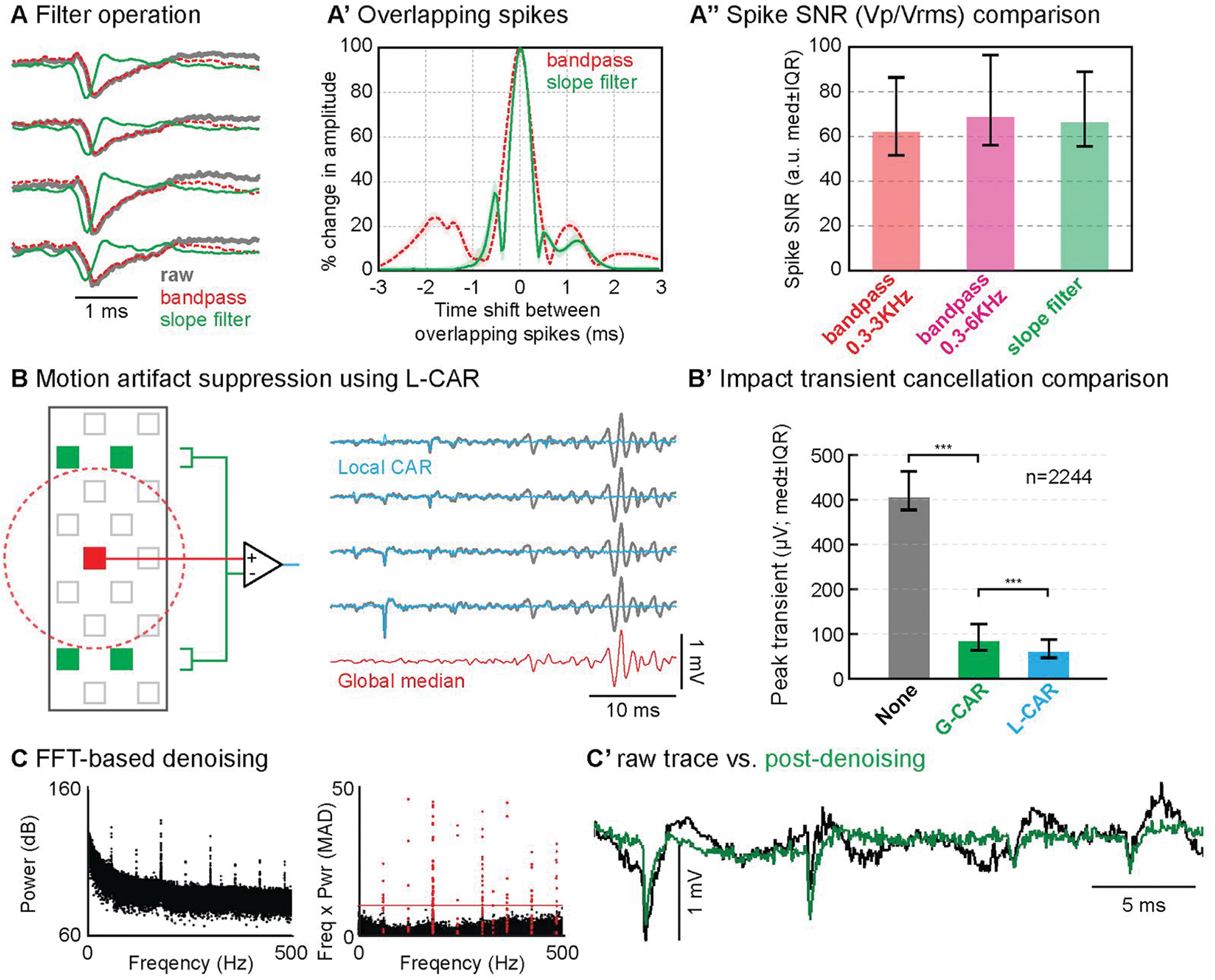
Signal preprocessing and denoising operations. (**A**) We apply a smoothing slope filter (Savitzky-Golay) to remove the LFP powers instead of the commonly used bandpass filter (Butterworth 300 - 6000 Hz) to reduce the occurrences of overlapping spikes. (**A’**) The slope filter exhibits a shorter sensitive time range between two overlapping spikes compared to the bandpass filter. (**A’’**) The slope filter results in a similar spike SNR (Peak spike amplitude / Vrms) compared to the bandpass filters. (**B**) Common average referencing (CAR) is used to remove external noise that commonly affects all channels. Local CAR (L-CAR) provides a better noise cancellation for spatially varying noise along the extended shank length. We computed the background noise for each site by averaging the neighboring sites, while excluding the immediately adjacent sites to preserve the spike waveforms. (**B’**) Significant reduction of the noise generated by physical impacts after applying the common average referencing. Physical impacts were applied to the head implants (12 events, 187 sites). L-CAR performed significantly better than G-CAR (two-sample T-test, *P*<0.001). (**C**) Narrow-band noise outliers are shown in the frequency domain (left), and these outliers are automatically detected after correcting for the 1/f trend by plotting the frequency-power product as a function of the frequency. Frequency-power products are expressed in the MAD (median absolute deviation) unit by dividing by the MAD values computed for each frequency bin (1 KHz width). Outliers exceeding 10 MAD (red line) are removed by setting their FFT coefficients to zero. (**C’**) Denoised signal (green trace) is obtained in the time domain by applying inverse-FFT to the cleared FFT coefficients.

Noise sources such as motion artifacts and background spikes overlap with the spiking signals in the frequency domain making them difficult to separate using filters alone. These noise sources are physically distant from the recording sites, and thus add highly correlated noise waveforms to a broad range of neighboring sites. Using this spatial redundancy, it is possible to remove the locally correlated noise from each site by averaging its neighboring sites. Since for high-density recordings spike waveforms spread to immediately adjacent sites, we calculate the local averages after excluding the nearest adjacent sites to prevent inadvertently subtracting relevant spike waveforms (Figure 2B, left). We refer to this strategy as local common average referencing (L-CAR) in contrast to the global common average referencing (G-CAR) [12], which subtracts the average of all sites from each site. L-CAR performs significantly better than G-CAR in cancelling the motion artifacts generated from physical impacts (Figure 2B’), since L-CAR captures the gradual spatial variation of the noise profile across the probe length. As will be discussed, the L-CAR operation is efficiently implemented in GPU and only adds an additional 10% of the total processing time.

Electrical noise from power lines and digital electronics often appears in neural recordings. These are characterized by multiple narrow-band peaks in the frequency domain (Figure 2C). Narrow-band noise as such typically undergoes dynamic changes such as transient appearance and disappearance of frequency peaks as animals move with respect to the location of noise sources [13]. Although notch filters can filter out manually specified frequency peaks, this is not adequate for dynamically varying noise with multiple noise peaks.

It is possible to automatically remove dynamically varying noise outliers in the frequency domain by applying Fast Fourier Transformation (FFT), setting the outlier coefficients to zero, and applying inverse FFT to obtain a denoised signal. Noisy outliers clearly stand out after detrending the power vs. frequency plot using the characteristic 1/f noise power in neural recordings (Figure 2C). To even out the variations of baseline power across frequency, we normalize the power-frequency products by dividing with the MAD (median absolute deviation) values computed in each frequency bin (1 KHz width). This FFT-based denoising method can adapt to changing noise conditions by updating the noise outlier detection in each time step (1 minute). This denoising step adds only additional 10% of the total processing time thanks to the GPU-based FFT operation.

### Feature extraction and clustering strategies suitable for HD probes

Extracellular probes with closely packed sites (20-30 μm center-to-center site spacing) typically pick up multiple spikes from the same neuron on adjacent sites [Anastassiou et al, 2015]. Accurate classification of these spiking events requires correct grouping of the redundant spikes spreading across adjacent sites. We perform the spiking event detection for HD probes in two steps. First, spikes are independently detected from each channel by searching for the turning points exceeding the detection threshold set automatically (6× MAD) [7]. Next, we assign the largest spike from each spiking event (center spike) by comparing each detected spike to its neighboring spikes (within 50 μm radius and ±0.5 ms) to determine the local maximum (Figure 3A). For each center spike, we extract the spike waveforms (1 ms) from a fixed number sites (n=10) surrounding the center site. In comparison, conventional event grouping strategy based on flood-filling operation yields highly variable event size (Figure 2A’), which is defined as the number of sites exceeding the event-grouping threshold (2×Vrms) [8,14]. In order to reduce the clustering errors caused by the variable event sizes, we extract a fixed-size spatiotemporal window surrounding the local maximum point (event center) where the SNR is the highest per spiking event.

**Figure 3.**
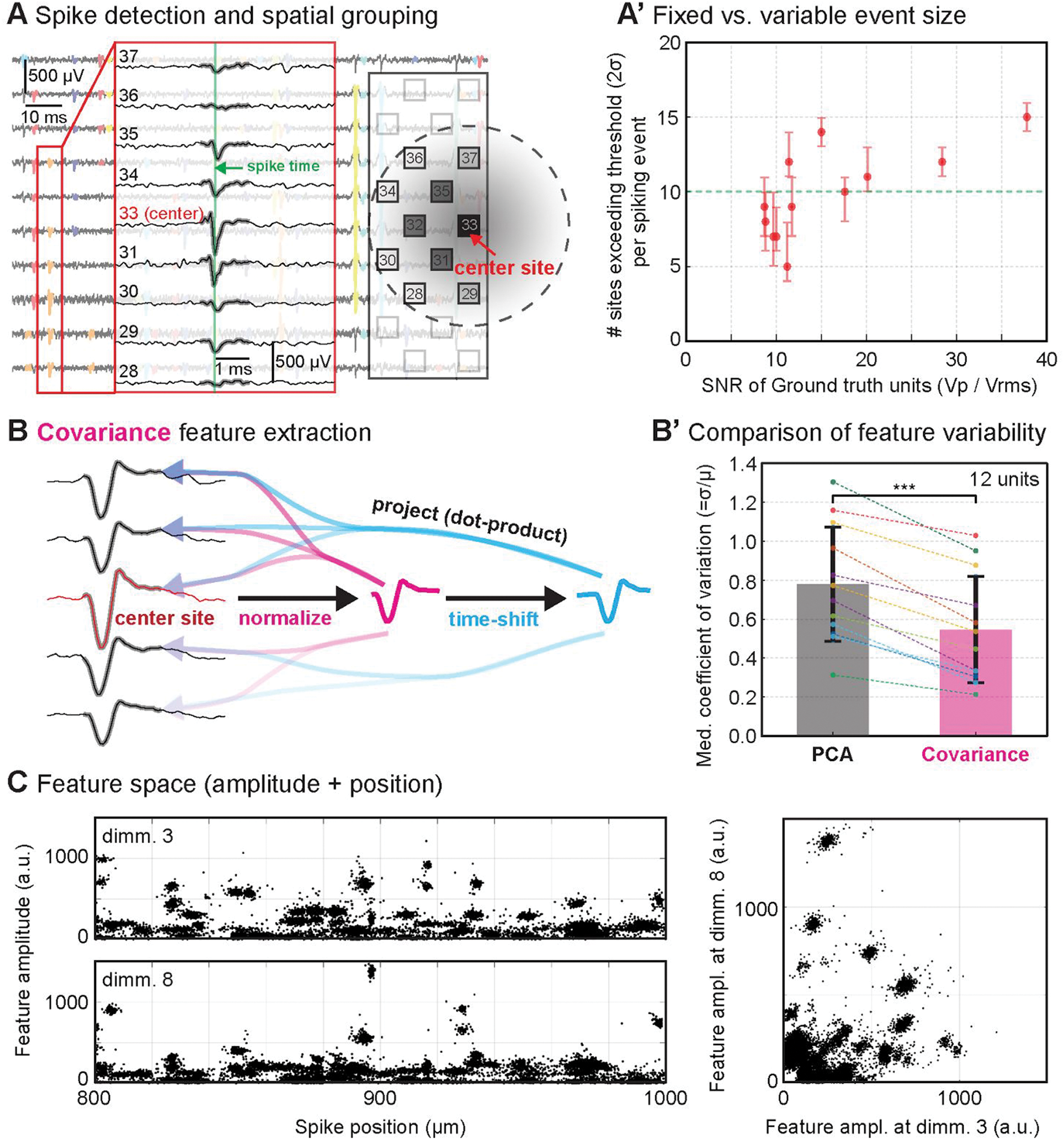
Spike grouping and feature extraction. (**A**) Single spiking event is detected by multiple adjacent sites. For each spiking event, we capture the spike waveforms from a fixed number of sites (n=10 within 50 μm) centered at the site containing the largest spike (center site). (**A’**) Spiking event size determined by the flood-filling algorithm generated a variable number of sites exceeding the threshold (2 SD). Instead, we used a fixed number of sites per event by taking spike waveforms from ten sites proximal to the maximum spiking site (green dashed line).(**B**) Instead of using the conventional PCA features, we used the covariance of spike waveforms between sites as features for clustering. For each spiking event, we computed the covariance between a spike waveform from each site and the template waveform (magenta trace), which is taken from the center site after normalizing. In order to account for the time delays between sites, we generate a second template waveform (cyan trace) by shifting the first template waveform forward in time (0.1 ms). (**B’**) Our covariance features exhibited significantly less variability per unit (*P*<0.001) compared to the PCA features. The variability per unit is quantified as the median Coefficient of Variation (CV) across 20 features per event (two features per site). This can be explained by the robustness of the covariance feature in the presence of independent noise between sites. (**C**) We estimate the each spike position along the probe length by computing the centroid, using the site locations weighted by the squared amplitudes. We then construct a single feature space for all spiking events from all sites by combining the spike positions and the feature amplitudes (20 dimensions = 10 sites/event × 2 features/site). Spikes are sorted in parallel by computing the distances to their immediate spatial neighbors (±5 μm) for efficient computation.

Spike-sorting requires the event waveforms to be represented as a set of low-dimensional features, so that a large population of spikes can be clustered efficiently. The dimensionality reduction step is typically performed using principal component analysis (PCA), which projects each spike waveform to a set of principal vectors to determine the PCA scores. PCA projection requires a common set of basis vectors to be computed from a population of spike waveforms. However, noise-induced temporal misalignments between the principal vectors and spike waveforms increase the feature variability. We introduce a robust feature-extraction strategy based on the channel-covariance that can tolerate spike timing jitters. The channel-covariance feature is computed by projecting spike waveforms to the normalized waveform from the center site (Figure 3B). Since the covariance calculation averages out the independent noise between channels, this method can extract robust features even from sub-threshold (4×Vrms) spike waveforms. Moreover, the spike timing shift does not affect the covariance features because the relative timing between neighboring waveforms remains constant. For a typical spiking event, spikes from two adjacent sites exhibit similar waveform shapes with a brief time delay (fraction of ms). Since this temporal relationship is a reliable feature of a given neuron, we capture this information by projecting spike waveforms to a time-delayed version (0.2 ms) of the center-spike waveform. Our covariance-based features exhibited significantly lower variability (*P*<0.001, n=12, paired T-test) than the PCA features (Figure 3B’), as quantified by the Coefficient of Variation (CV) from hybrid ground-truth units (see Methods).

In addition to the covariance between sites, we also include estimated spike positions in our feature set. The spike positions can be estimated using multiple, coincident spikes appearing on closely-spaced sites on a HD probe. We estimate the spike positions using the center-of-mass calculation by averaging the site coordinates weighted by their spike amplitudes (Figure 3A,C) [15]. Such additional spatial information improves the sorting accuracy, and allows faster clustering in parallel by spatially dividing the population of spikes. The spike positions also provide means to correct for the probe drift as will be discussed later.

Conventional spike-sorting methods employ a clustering algorithm based on fitting a mixture of Gaussian distributions (GMM: Gaussian Mixture Model) whose parameters are determined by the Expectation Maximization (EM) algorithm [8,14,16]. However, this approach is not appropriate for HD probes since the algorithm does not scale well for the increasing number of sites (see Methods). Furthermore, the assumption of Gaussian distribution does not hold well for real recordings exhibiting bursting dynamics and probe drifts. Instead, we apply a state-of-art clustering based on fast search of density-peaks (DPCLUS) [17]. This simple yet powerful clustering algorithm requires only a minimal set of parameters which works robustly without requiring adjustments. DPCLUS is agnostic to the underlying distribution shapes and can automatically discover the number of clusters by identifying the local density peaks. The clustering operation is performed in GPU to rapidly compute the pairwise distances between spiking events, and requires an insignificant fraction (<5%) of the total processing time (Figure 7B,C).

### Spike sorting validation using multiple ground truth

It is critical to validate the quality of spike sorting across a spectrum of ground-truth datasets where spike timing from different neurons/units are known. Our choice of algorithms and their parameters was guided by their overall performance on a large collection of such datasets. The neuroscience community has recently generated multiple types of ground-truth datasets for HD probes. Ground-truth datasets provide a subset of spikes with known neuronal identities, which permit a direct quantification of spike sorting accuracy. There are multiple methods of generating ground-truth information, each with strengths and limitations. In order to independently assess spike sorting quality of JRCLUST, we combined three types of ground-truth datasets generated by 1) paired intra/extracellular recordings, 2) manual curations followed by random shuffling, 3) biophysically detailed simulations of a large neural population.

First, we used simultaneous intracellular and extracellular recordings from same neurons in slice preparations ([10,18]) and in vivo ([10,18]; Figure 4A). Although this approach (paired recording) is limited to one known neuron per recording session, it provides precise timing of all extracellular spikes emitted by a single neuron recorded via intracellular whole-cell or juxtacellular patching. Second, we selected a set of manually well-isolated units, and injected their spikes at randomly chosen sites and times (Figure 4B). We refer to this as a “hybrid ground-truth” [9,19] since we automatically assigned spike timing and positions after manually isolating them. Although hybrid ground-truth datasets provide multiple known units per recording session, this method is limited to testing high SNR units that are well isolated. Third, we simulated a biophysically realistic network of neurons, with each neuron reconstructed with realistic morphology and detailed individual channel dynamics. The simulation generated extracellular voltages at each virtual recording site that mimics the actual site layout, and intracellular voltages for all neurons in the simulated tissue volume surrounding the virtual probe (Figure 4C). Our *in-silico* ground-truth dataset provides the neuronal identities for the entire extracellular spikes being generated, thus allowing a parametric study of the spike sorting accuracy across a wide range of SNR, and for different site layouts.

**Figure 4.**
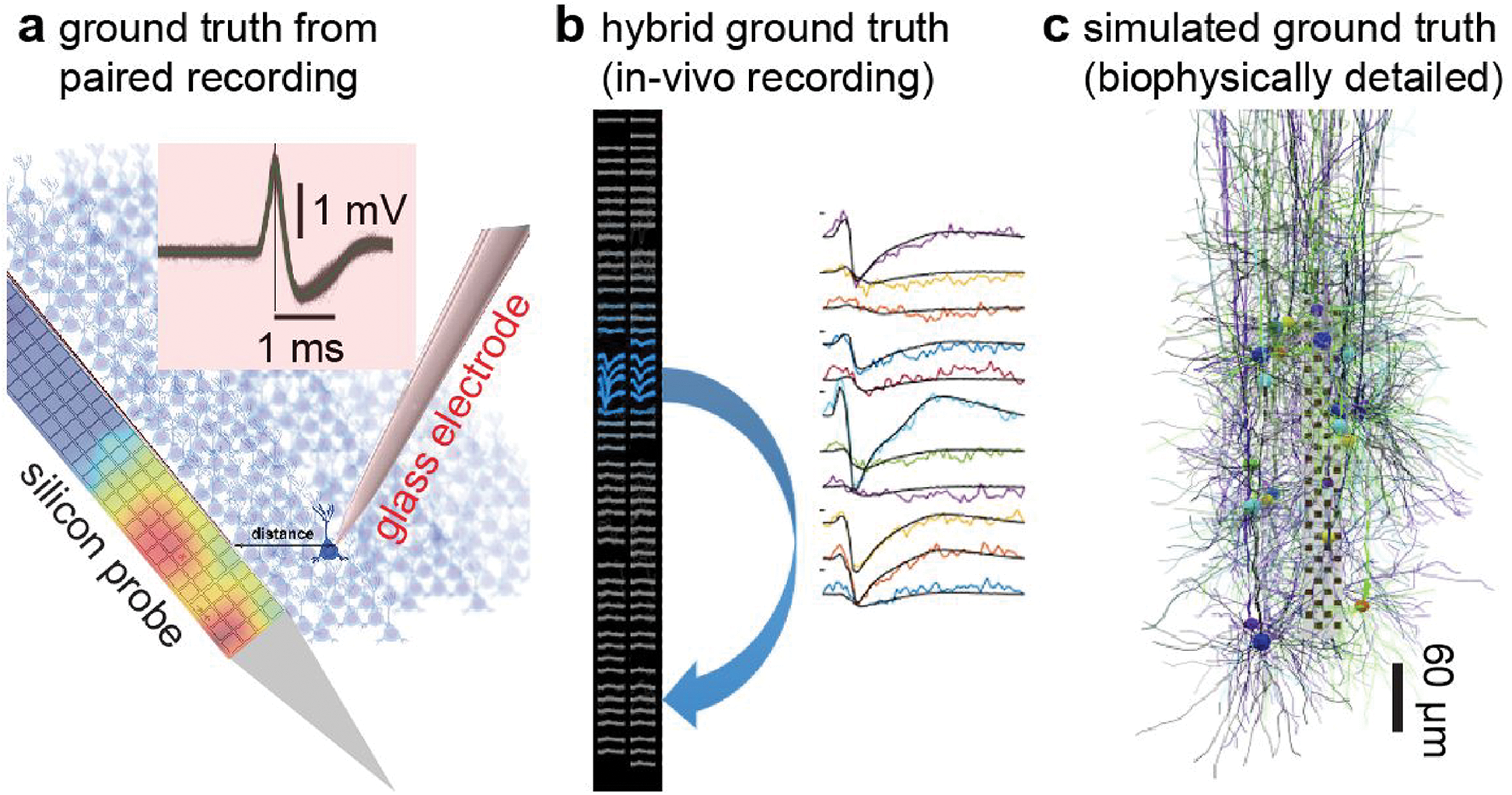
Three types of ground truth used for validation of spike sorting accuracy. (**A**) Paired recording is obtained by positioning a glass electrode near (50-100 μm) a silicon probe. The glass electrode (intra- or juxtacellular recording) provides a precise spike timing from a single neuron. (**B**) It is possible to synthesize a ground truth unit by manually isolating a well-separated unit, and adding its spike waveforms to an arbitrary location at random times. Since it combines manual and automated procedures to generate a unit, this is referred to as a “hybrid” ground truth. (**C**) Large neuronal population (~1000) surrounding the probe can be simulated in a biophysically detailed manner. The simulation captures individual channel dynamics from multiple compartments using realistic neuronal morphologies. The simulated ground truth provides complete intracellular voltage information for all neurons being simulated.

We quantified the accuracy of spike sorting using two metrics, false positives and false negatives, based on the fraction of spikes shared between ground-truth units and matching sorted units (Figure 5A). We first find the best-matching sorted unit for each ground-truth unit that shares the greatest number of spikes, and quantify the fractions of mislabelled or entirely missed spikes. The false-positive rate (*FP*) is defined as the fraction of spikes from a sorted unit which are incorrectly assigned to other ground-truth units (*FP=(n1-n3)/n1*; *n1*: number of spikes in the ground truth unit; *n3*: number of spikes commonly found in the ground-truth unit and the best-matching sorted unit). Likewise, the false-negative rate (*FN*) is defined as the fraction of spikes from a ground-truth unit while not present in the matching sorted unit (*FN=(n2-n3)/n2*; *n2*: number of spikes in the best-matching sorted unit).

**Figure 5.**
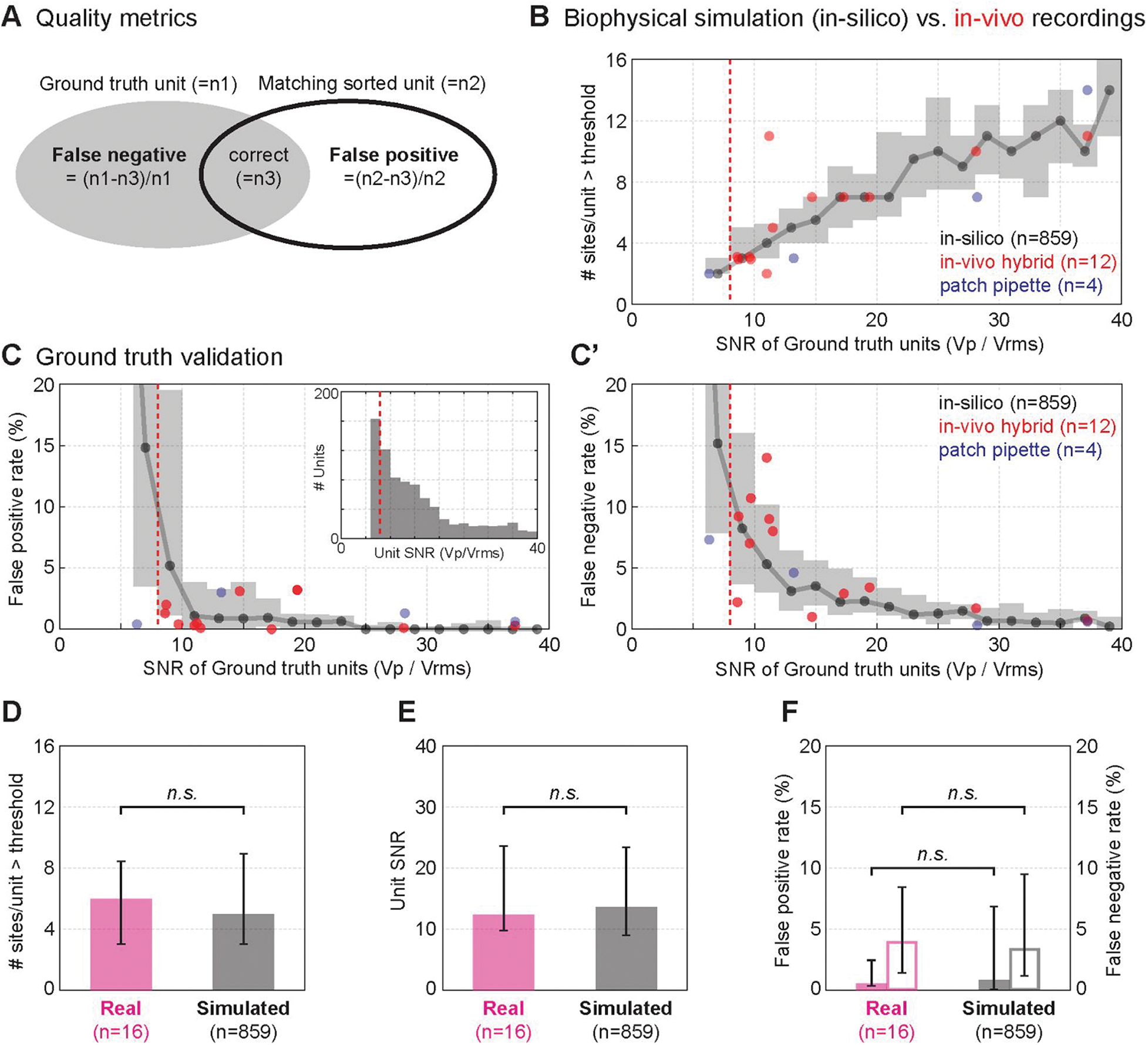
Ground truth validation of our spike sorting pipeline. (**A**) We quantified the accuracy of our spike sorting outcome by measuring the fraction of false positive and false negative spikes per unit with a known identity (ground truth unit). (**B**) All three types of ground truth datasets exhibited similarly increasing spatial spread as a function of the normalized spike amplitude (SNR=Vp/Vrms). The probe site spacing for all datasets falls within 20-40 μm. The shaded area and the black dots indicate the interquartile range and median values for the simulated ground truth (*in-silico*) throughout this figure panel. (**C**) Percentages of the false-positive and false-negative spikes are shown for each ground truth unit as a function of the normalized spike amplitude (unit SNR). Spike-sorting errors increased as the unit SNR decreased, and the median errors exceeded 10% for units SNR below 8 as indicated by the dashed red lines. We found a greater number of units having a lower SNR as expected by the inverse relationship between the spike amplitude and the distance to a neuron (inset). (**D-F**) No statistically significant differences (two-sample T-test) are found between the real (magenta) and simulated (black) ground-truth units in terms of the number of sites per unit exceeding the detection threshold (4 Vrms) (**D**), unit SNR (**E**), and false positive and negative rates (**F**). Median values and interquartile ranges are shown for the ground-truth units exceeding 6 SNR.

We found a close match amongst all three types of ground-truth datasets (Figure 5B,D,E) recorded from similar site layouts (see Methods). As the amplitudes increased, a higher number of sites contained spikes above the detection threshold (Figure 5B). The quantitative relationship between the spike amplitudes and the number of sites exceeding the detection threshold agreed between the real (patch pipette and hybrid *in-vivo*) and simulated (*in-silico*) recordings, and no significant differences were found between the real and simulated recordings in terms of the number of sites per unit and the unit amplitudes (Figure 5D,E). In order to account for the variations in the baseline noise levels, we express the spike amplitudes in SNR units for a fair comparison across multiple datasets by normalizing with the baseline noise (RMS voltage after correcting for spikes).

We characterize the spike sorting accuracy as a function of the SNR of averaged unit waveforms at the event-center sites (unit SNR). There was a strong dependence of the false positives and negatives on the unit SNR, and this trend closely agreed between the real and simulated recordings (Figure 5C). For the units with amplitudes above 6×Vrms (SNR>6), the median sorting errors were under 5% (Figure 5F), and the error rapidly increased as the unit SNR fell below 8. The sortings errors were not significantly different between the real and simulated ground truth datasets (Figure 5F). These results suggest that it is necessary to impose a criterion based on the unit SNR to ensure the quality of spike sorting. With this framework, these quality metrics enable the discovery of more accurate sorting methods that can yield more usable units at a lower range of SNR. The *in-silico* ground-truth captured an increasing number of units having lower SNR, as expected by the increasing population of neurons further away from the electrode (Figure 5C, inset).

### Effective site layout patterns for high sorting accuracy and unit yield

In order to determine the most efficient site layout to improve the spike sorting outcome, we assessed both *unit yield* and *sorting quality*, which are considered to be the two most important features to be optimized. We define the unit yield as the number of ground-truth units exceeding 8 SNR, and the sorting quality is measured by the false positive and negative rates. We selected four probe layouts with the identical number of sites and vertical span to test that were under consideration for wide-scale production by the Neuropix Consortium. We simulated these four site layout patterns using the same computational framework described in Fig. 4A (see also Methods) under identical conditions to assess their effectiveness. This side-by-side comparison would not have been possible in a real experimental setting.

The four site layout patterns each contained 60 sites, and identical 20 μm vertical spans, but the horizontal site spacing varied between 16-48 μm (Figure 6A). Generally, as the horizontal spacing increased the quality of the spike sorting decreased (Figure 6B), however inversely the unit yield increased (Figure 6C). Interestingly, by using a staggered 4 column (*4col*) site layout pattern we produced one of the highest unit yields with the highest spike sorting quality. This can be explained by the fact that the site spacing matched the spatial scale of the signal spread in extracellular space by offering an optimal combination of spatial redundancy through a uniform site spacing distribution in both horizontal and vertical dimensions.

**Figure 6.**
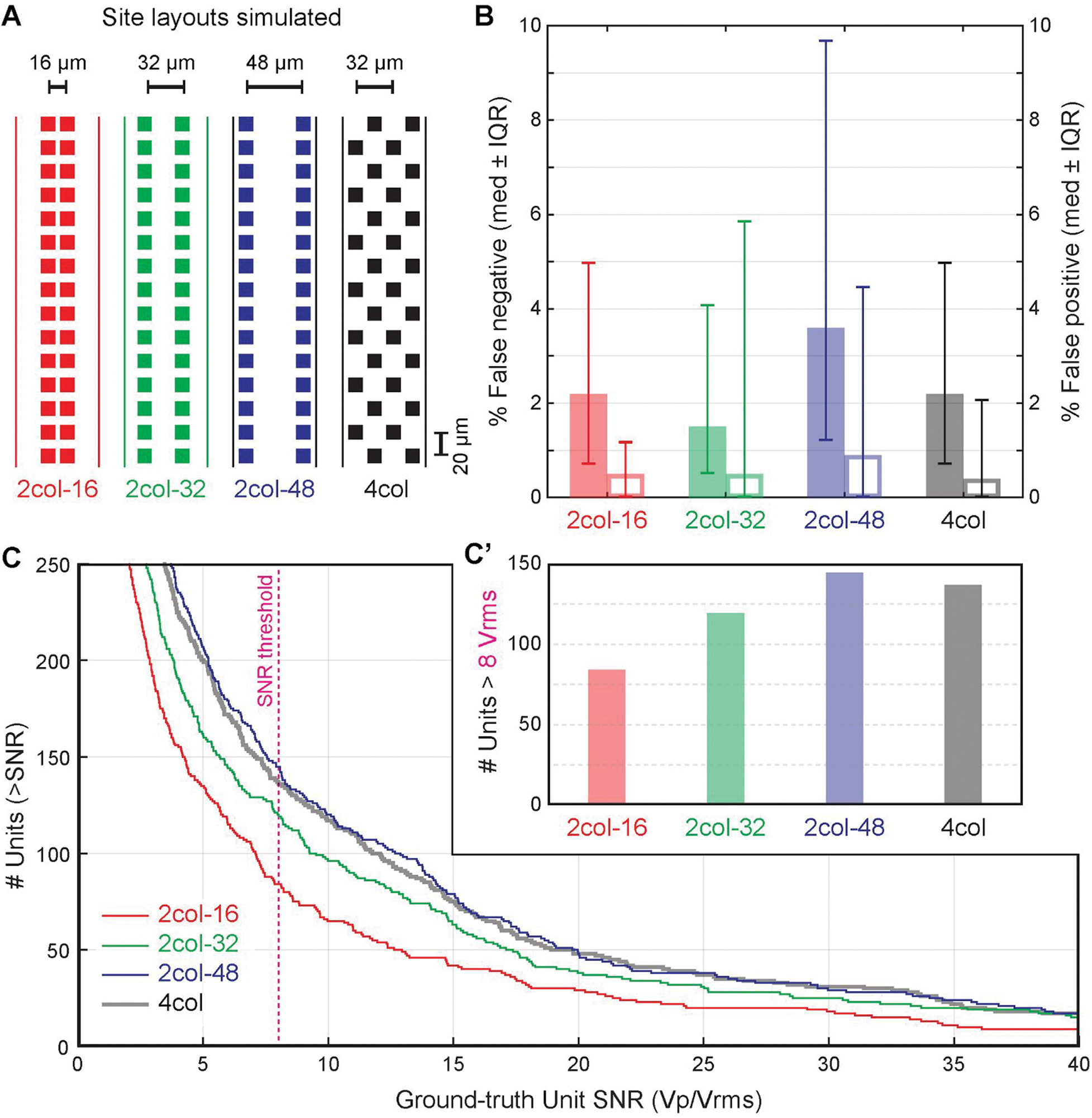
Comparison of site layouts on sorting quality and unit yield.

(**A**) We simulated four site layout patterns containing 60 sites each. The horizontal site spacing varied between 16-48 μm but the vertical spacing was fixed at 20 μm. (**B**) The sorting accuracy generally decreased as the horizontal spacing increased. The median values and interquartile ranges are shown, and solid/open bars indicate the false negative/positive rates. The four-column layout (*4col*) with a staggered pattern had an intermediate accuracy between the 32 μm and 48 μm spacing. (**C**) The unit yield was the lowest for the narrowest layout pattern at 16 μm while wider horizontal separation increased the number of units picked up by the probe. *4col* layout offered unit yields virtually equivalent to *2col* 48 μm spacing while offering noticeably better spike sorting quality. (**C’**) The unit yields, defined as the number of units exceeding 8 SNR, are shown for the four simulated site layout patterns.

### Fast and scalable performance of our spike sorting pipeline

With the massive amount of data collected from HD probes the analysis processing time much catch up to allow efficient interpretation. This is especially important given the increasing number of sites on probes in development. Our analysis pipeline achieved high speed and scalable performance without sacrificing the quality of analysis.

Figure 7A shows three example recordings from a high density probe with 120-sites. The total processing run-time speed of our complete spike sorting pipeline is ~10x faster than real time recording speed from a 120-channel probe using a single workstation equipped with a GPU. Because processing time scales linearly to the number of channels, this translates roughly to the processing speed of a 1000 channel probe recording at real-time allowing analysis time to match recording time enabling real-time results.

We found that the processing stage is uniformly divided across the entire processing pipeline demonstrating a balanced optimization with no major bottleneck stage (Figure 7B). The analysis scales linearly to the recording size for the preprocessing stage due to its strong dependency on the data I/O speed. The rest of the process scales linearly to the number of spikes as these operations are mostly computation speed limited. Traditionally, the spike clustering stage is the major processing bottleneck, however GPU implementation of our algorithms offered significant speed enhancements (65x) over CPU. Additionally, the GPU computation significantly boosted the overall processing speed of the other stages (Figure 7C).

**Figure 7.**
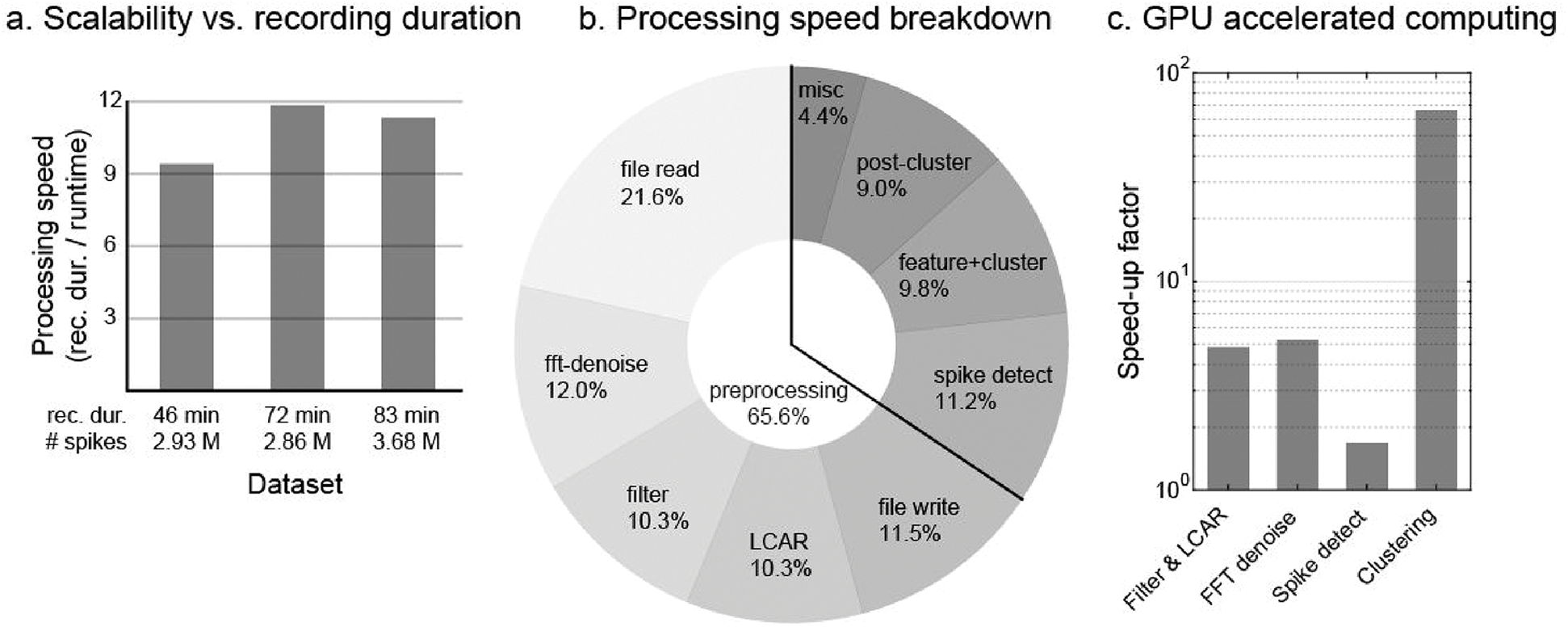
Speed performance. (**A**) The total processing run-time speed of our complete spike sorting pipeline is ~10x faster than real time recording speed from a 120-channel probe using a single workstation equipped with a GPU. (**B**) The breakdown of the processing time by each stage is shown here. The processing stage is uniformly divided across the entire processing pipeline. (**C**) GPU implementation of our algorithms offered significant speed enhancements over CPU. In particular, the clustering algorithm showed the largest speed improvement of 65 times, which typically has been the slowest step of the entire processing pipeline.

### Automated probe drift detection and correction

Probe drift is a common problem when recording with behaving animals and often causes spike sorting errors due to a non-stationary signal. This introduces spike amplitude fluctuations of the same unit in neighboring sites caused by the tissue movement bringing sites closer or further away from the spiking source. These continuously drifting amplitudes can cause erroneous cluster splitting (Figure 8A) or the loss of tracked units. However, an HD probe can potentially solve this challenge by offering many additional sites along its depth and enabling experimentalists to track these moving units.

**Figure 8.**
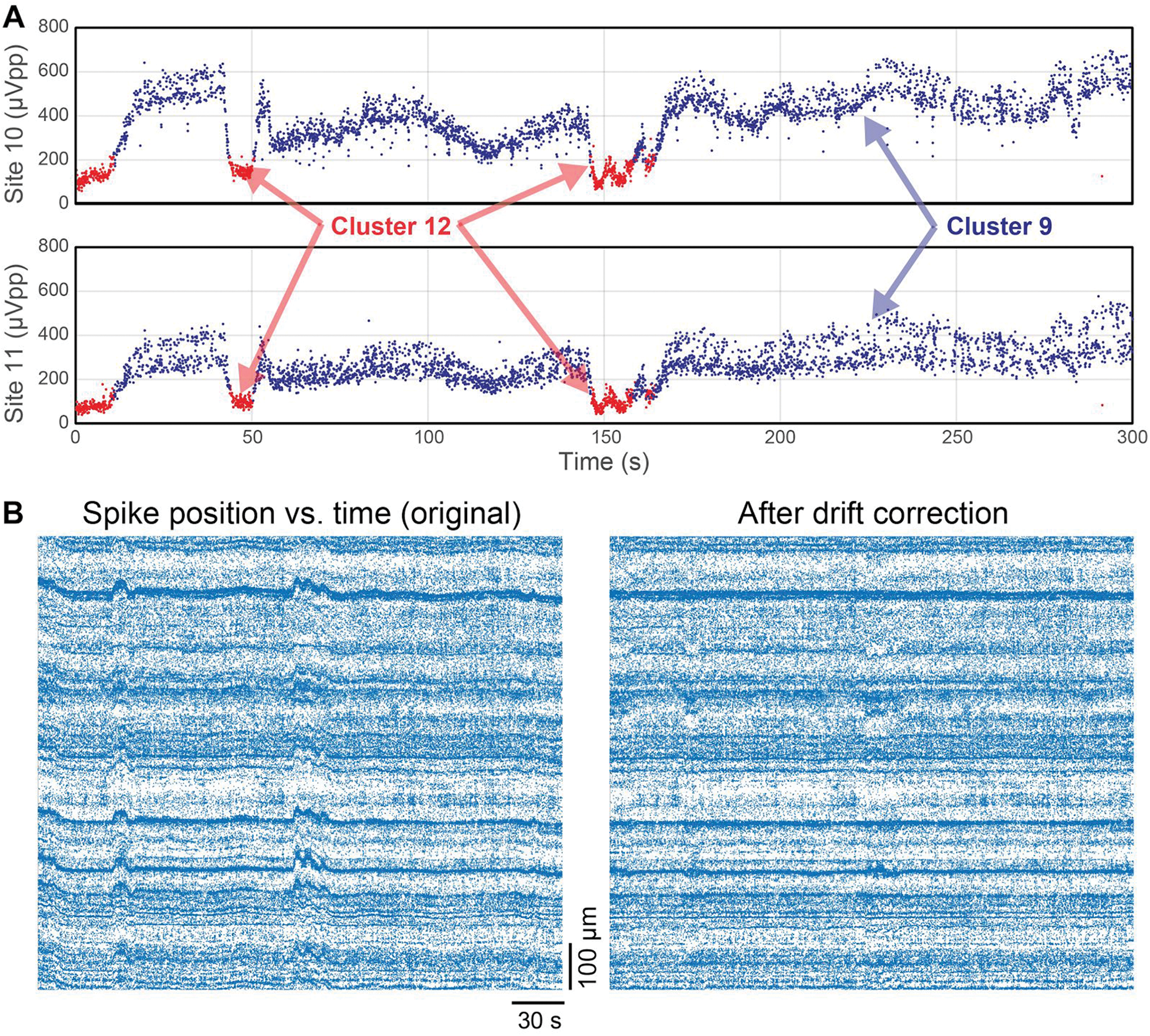
Probe drift detection and correction. (**A**) Probe drift across time manifests itself by co-modulating the spike amplitudes of the same unit in neighboring sites. This is due to the tissue movement bringing sites closer or further away from the spiking source and is exhibited by fluctuating spike amplitude over time. This fluctuation occurs concurrently across adjacent sites. The drifting spike amplitude causes erroneous cluster splitting. (**B**) The global tissue drift across time is apparent across the whole shank when spike positions are plotted across time. This visualization (left) shows bands of activities that fluctuate coherently across the depth of the tissue with each plotted point representing an individual spiking event. We corrected for this global drift across time by estimating the probe position by using cross-correlation of the spike positions. The corrected spike positions (right) show constant activity band positions across time.

Using the already computed spike positions, as a function of time, allows us to identify global probe movements (Figure 8B, left) by revealing activity band patterns that fluctuate coherently over time. We correct the spike position by subtracting the probe position as computed at each time bin (1 s) to result in constant activity band positions (Figure 8B, right). We estimate the probe position by computing the cross-correlation of the spatial profile to determine the optimal vertical spatial shift between neighboring time bins offset the drift. This step is performed in *JRCLUST* before the clustering stage.

## Discussion

### Summary statement and significance

Fast, accurate and robust spike sorting is key to bridging the existing gap between the capabilities of current recording technology to gather massive amounts of neural data and the software needed to interpret that data in a scalable way. Up to now, a bottleneck has existed due to the need to manually identify clusters of spikes by drawing boundaries by hand. The impact of the new *JRCLUST* is that there is now a scalable, proven, automated solution to spike sorting that can tolerate real recording conditions with noise, probe drift, and motion artifacts from behaving animals. While there exist other clustering algorithms that are applicable to specific datasets after extensive parameter tuning, *JRCLUST* can handle a wide range of datasets using a pre-optimized parameter set which makes it practical for wider use. Also our processing speed and modular approach allows for rapid cycle innovation and practical pathways to interpret massive volume of extracellular recordings. This advancement sets the foundation for future innovations in algorithmic performance, the use of real and simulated ground truth datasets, and probe hardware optimization. Our platform can also potentially be extended to become a community resource for standardizing algorithm validation using common performance metrics and datasets.

### New spike sorting approaches enabled by improved extracellular electrodes

Since the launch of the BRAIN initiative in the U.S., Human Brain Project in Europe, and the Brain and Mind project in Japan there have been major coordinated efforts to advance neural probe technology. Thus far these advances have been rapidly accelerating in these areas: 1) higher site density, 2) extended spatial coverage, 3) high fidelity recording, and 4) integration with cell-type specific stimulation tools (Ed Boyden, Neuropixels paper, Neuroseeker Adam Kampff paper, microLED by Fan Wu). These long-awaited methods for better recording bring the field within reach of a major ambition: whole brain recording at a single neuron, single spike resolution. However, parallel advancements in software are necessary to achieve this aim. To take full advantage of these hardware capabilities, new approaches to interpretation, automated quality controls, as well as solutions for analyzing a massive amount of collected data need to be developed.

Access to industry-grade semiconductor foundries have allowed for the creation of probes with closely packed small recording sites and multilayer traces. These provide a higher resolution that can detect activity in small cells (under 10 um) and fine axonal features. This level of resolution allows us to distinguish cell types through the subtle differences in spatial pattern of the electrical field. While critics of this approach have worried that the amount of data produced by this high resolution would be time consuming and burdensome to manually curate, and thus un-scalable, the increased information provided by these additional sites allows us to perform very accurate automate spike sorting through a algorithm development. *JRCLUST* has been designed to capture fine spatial patterns and features from these closely packed sites allowing for the reliable discrimination between different neurons (Figure 3A,B).

Whole brain depth probes, such as the Neuropixels probe, allow for increased spatial coverage spanning many different brain areas. Previous clustering algorithms could not scale to handle recordings from extended spatial coverage without efficient spatial division strategies. However, *JRCLUST* makes better use of the spatial information of neural activities to enable a seamless divide and conquer strategy, taking advantage of parallel computing algorithms and hardware such as GPU and multiple CPU cores. Because these probes do not require physical translation, it is now possible to chronically record from a fixed location for many months opening pathways to long-term cell activity tracking studies. A fundamental challenge to this is brain tissue movement that introduces non-stationary activity signatures obfuscating linked cell activity between recording sessions. *JRCLUST* took advantage of extended vertical site span to automatically correct tissue drift by tracking fine probe movements using correlated global shifts in activity profile.

The uniformity of low impedance recording sites and low noise recording electronics opens up the possibility of a solution for global noise cancellation. Prior to the advent of integrated recording electronics, external noise used to cause variable responses at each recording channel because of variable signal paths to the amplifiers. On-chip integration of amplifiers and digitizers has mitigated this issue by minimizing the length of the signal path from detection to recording leading to a greater homogeneity of each site’s response to external noise. Taking advantage of this we created an effective noise cancellation algorithm for external noise sources that commonly occur during behaving animal experiments (Figure 2B,C).

Concurrently, there are advances in stimulation technology that give researchers the opportunity to influence genetically targeted specific cell types as well as record neural activity. This is made possible by the integration of electrical and optical components in one probe [20–24]. However, unlike optical recordings that provide cell type information, up to now the weakness of extracellular recordings has been a lack of exact cell type information. There are now efforts to relate electrical activity patterns that are specific to genetically identifiable cell populations [25–27]. *JRCLUST’s* ability to cluster unit activities could be extended to identify specific cell types through use of such devices and the generation of ground truth data inclusive of cell type metadata. Another possible approach to generate such data is to combine electrical and optical recordings using two-photon microscopy [28].

### Quality enhancements through simulated and real ground truth

There are currently no established methods or concerted efforts to develop quality standards for comparing the efficacy of spike sorting approaches. Establishing such standards is important to bringing the field’s parallel efforts in algorithm development and ground truth data generation together. To that end, the provision of long-term professional support for the community could be a sustainable strategy for the discovery of better algorithms along with new approaches to the aggregation of simulated and real ground truth.

*JRCLUST* lays the foundation to meet the community’s needs for spike sorting by expanding support for popular algorithms unified under a common quality validation and interactive data visualization platform. We already have integrated community-contributed ground-truth datasets into our quality validation and comparison tools. Also, *JRCLUST* supports a wide range of extracellular probes and recording hardware from multiple vendors. Comprehensive documentation and tutorials will further be developed and posted online so that users can quickly learn and add custom functionalities. *JRCLUST* will be continuously improved as the community aggregates new ground truth data, both simulated and real, for analysis and comparison.

Simulated neural activity allows for the rapid exploration of different recording conditions through the manipulation of parameters (such as SNR, cell types, and firing rate) that influence spike sorting quality. Through this simulated platform we can quickly discover site layout patterns optimized for specific brain regions and test the efficacy of future spike sorting algorithms. It also allows for the rapid exploration of optimal approaches to specific brain regions with very different spatial organization and structures. For example, the hippocampal CA1 consists of a layer of closely packed pyramidal cells which may require a different recording and analysis method than the striatum which contains a more dispersed population of medium spiny neurons. Although our simulation is based on one of the most detailed biophysical models of neurons at the individual channel level, real ground truth studies are still necessary to build better-calibrated simulations.

Although real ground truth data sets, which require access to intracellular voltages, are difficult to obtain and limited in scale, there are technological advances that can make such aggregation less burdensome. For example, researchers at HHMI-Janelia Research Campus have developed electrodes that can simultaneously measure from both inside and outside the cell membrane in a single device and multi-cell intracellular recordings can assist in the creation of high yield ground truth. Further, there is an intracellular electrode positioning system being developed that can easily target a single cell [29] by triangulating current pulses. All of these advances can simplify the collection of this critical validation data.

Two-photon calcium imaging combined with electrical recording within the same detection volume could provide a scalable way to obtain intracellular measurement by building on the combined strengths of each method: optical identification of genetically targeted population, detailed structural imaging and high temporal resolution. Alternatively, integrated optical probes (Fan Wu and IMEC Hoffman) are available that can excite specific cell types that are expressing opsins, providing a pathway to generate cell-type specific ground truth. These are exciting opportunities for cell-type identification and spatial information about cells that can be used for spike sorting validation.

### Support real-time feedback experiments by integrating data acquisition system and hardware-accelerated computations

Real-time analyses of brain recordings are necessary for exploring causal relationships between neural dynamics and sensory-motor observables by providing closed-loop feedback. However, previous approaches to spike sorting were too slow to be useful for real-time interaction with the brain because the processing load could not be efficiently distributed in a local parallel computer architecture. Networked parallel computing is also not adequate to the task because of data transmission delay and latency over such a network due to the massive volume of sensor data. To address this problem, we used an efficient GPU implementation and multi-CPU core architecture in a local computing environment. We also performed data reduction through event feature extraction and effectively used on-chip cache memory to minimize unnecessary I/O operations. Additionally, we used a performance profiler to identify and improve the slowest and most memory hungry stage of the pipeline.

The experimental study of synaptic plasticity requires activity-based feedback at the synaptic integration time scale (<2 ms). While the algorithm as designed is theoretically capable of real time analysis it has not yet been implemented for online processing. To bring this process online, tight integration with data acquisition hardware is necessary to receive data as it is recorded. Also needed is the ability to provide real-time visualization feedback to users. Currently, most data acquisition hardware is reliant upon a USB interface, which cannot provide sufficient data bandwidth or reliable low latency for online processing. One remedy for this would be to migrate is a PCI-e interface as it provides roughly 4-8 GB/s. Direct data streaming to an FPGA board with a PCI-e interface could allow fast access to local computer resources through a direct memory access (DMA) interface [30]. Additionally, multiple GPUs and FPGA-based processing can further augment computational power.

### Cloud-based computing and data sharing

A web-based data hosting and analysis platform is essential to building a stronger neuroscience community by facilitating data and code sharing. By promoting collaborative behaviors, it will ultimately improve the transparency, reproducibility and efficiency of our scientific endeavors. Such a platform will let researchers upload in a compressed data format, enabling others to search and re-analyze the data on the server without having to download original datasets. This collaborative platform would encourage experimentalists and theorists to coordinate data collection and analysis efforts, thus ultimately accelerating the discovery of fundamental principles of the brain. The challenges facing this endeavor include different data formats, approaches to data analysis, and massive data that need to be shared.

To solve these constraints it is key to support data standardization efforts such as Neural Data without Borders (NWB), which is based on a widely used scientific format called the hierarchical data format (HDF5). Additionally, multiple data analysis methods need to be implemented on a common web platform, which would allow server-side computation without the need to download data for local processing. Further, raw data sharing demands too much bandwidth and storage, and sharing could better be facilitated by uploading spiking event information only which would dramatically reduce transferred data volume.

## Materials and methods

### Spike sorting pipeline

Here we describe key operations performed at each stage of our spike-sorting pipeline. We have developed a set of comprehensive and practical tools to address challenges that are often encountered in experimental settings. These include conditions such as electrical noise, motion artifacts and probe drift. We have taken a data-driven approach to develop our pipeline using ground truth datasets from real experiments and biophysically-detailed simulations. Our design choices were also guided by the simplicity and robustness of the algorithm under various dataset without the requirement of extensive parameter optimization. Finally, we implemented these simple yet powerful algorithms as efficiently as possible using parallel computing environment such as GPU and multi-core CPU. Our spike-sorting pipeline (Figure 1B) achieved robust performance for various datasets and real-time processing speed for HD probes (Figure 1A) using a single workstation equipped with a GPU.

### Filtering and denoising operation

Raw recordings contain spike waveforms lasting 1-2 msec and slower oscillations from the local field potential (LFP) and noise. The preprocessing stage prepares the raw traces for spike detection by allowing spike waveforms to pass while suppressing all other signals and noise sources. A high-pass filter then removes the slower oscillations such that spikes can be detected when the signal crosses a fixed threshold. Certain high-frequency noise sources require additional processing since they overlap with the spectral range of the spike signal. To accommodate this, notch filters can selectively block a specific range of frequencies, and they are effective at attenuating narrow-band noise sources such as those from power lines and electronic clocks (Figure 2C).

Neural recordings often exhibit transient noise sources with multiple noise peaks, which typically occur when animals move relative to the noise-generating sources. Such recording conditions with dynamic and complex noise components require more than a static notch filter with manual frequency tuning. Adaptive filters can effectively cancel these time-varying noise components by identifying and suppressing frequency outliers. Neural signals contain smoothly varying powers that are inversely proportional to the frequency and can be distinguished from the external sources exhibiting narrow peaks such as power lines and crystal oscillators. We detect such outliers in the frequency domain whose power significantly deviates from the expected (1/f) trend, and for each time step (~60 s), we apply FFT (Fast Fourier Transformation) on the raw traces to identify these outliers in the power-frequency product as a function of frequency (Figure 2C). The power-frequency products is normalized by the MAD (median absolute difference) score for each frequency bin (1 KHz width). The noise peaks are then detected by applying a threshold (10× MAD, median absolute difference), and removed by setting their FFT coefficients to zero. Denoised version of the signal in the time domain is obtained by applying an inverse FFT. This FFT-based denoising operation is performed in GPU, and it only adds about 10% to the whole processing time (Figure 7B).

Butterworth filter, which emulates traditional analog filter electronics, is one of the most widely used types of digital filter for spike sorting. However, infinite impulse response (IIR) filters such as the Butterworth filter introduce phase distortions and time delays. While the phase distortions can be corrected by introducing a second filtering operation in the time-reverse direction, this acausal operation adds a significant processing time delay rendering IIR filters unsuitable for real-time implementation. Furthermore, an IIR filter operation requires the data to be represented as a real number (floating precision), which requires the integer data (fixed precision) obtained from analog-to-digital converters (ADC) to be converted to the floating-precision format. This data conversion step introduces extra delays and requires 2-4 times greater memory. For high-channel count probes with hundreds of channels, the conventional filtering operation leads to an excessive computational overhead.

In comparison, our approach uses a significantly faster and more effective filtering operation for spike detection and sorting that only introduces a sub-millisecond time delay. Our filter implementation directly operates on the raw data format (16-bit integer) without requiring data type conversion, and further speed gain is realized by running on GPU. We applied a smoothed differentiation filter, whose coefficients are analytically derived from a least squares quadratic fitting and differentiation (Savitzky-Golay filter of order 1, 9 taps). The smoothing operation in our filter limits the amplification of high-frequency noise associated with a differentiation operation. The resulting waveform is well suited for spike detection since the differentiation operation removes the slow oscillatory component, and the negative slope of the initial phase of action potential is amplified (Figure 2A). Further, the differentiated spike waveform becomes narrower than the original waveform (~2 ms width) thus allowing a use of narrower time window (1 ms) to reduce the effect of temporally overlapping spikes (Figure 2A’).

### Common average referencing to remove background spikes and motion artifacts

Raw signals from two neighboring sites share correlated noise sources from external or internal to the brain. External noise sources such as the power line noise and the motion-artifacts from impacts or feeding have global effects on all recording sites. Many of these noise sources contain high-frequency components that overlap with the signal band. Common averaging referencing (CAR) scheme [12] can be effective at reducing the correlated noise by subtracting the averages across all channels from each channel. However, this strategy fails when the noise waveforms vary across sites, which is the case for HD probes containing hundreds of sites that extends several millimeters in length. For example, chewing or physical impacts typically results in noise waveforms that gradually vary across space with time delays. Global averaging fails to suppress such noise waveforms when there exists sufficient variations in waveforms and amplitudes across space.

In addition to the external noise sources, neural recordings contain background spiking activities from distant sources that are too weak to resolve but still contribute to noise. Spikes from distant neurons are added to a broad range of sites and they corrupt spike waveforms from nearby sources [10]. Spatially varying noise such as the background spikes and the motion artifact can be cancelled by applying a local common average referencing (L-CAR) scheme (Figure 2B,B’). Instead of subtracting the averages across all channels, each channel can be subtracted by the averages of its k-nearest neighboring channels. In order to prevent subtracting the spike waveforms from the nearest neighbors, we compute the local averages after excluding the immediately adjacent sites. For Neuropixels probe, we averaged four nearest sites after excluding the self and five nearest sites. The L-CAR operation was very effective at cancelling the correlated noise, impact transients and improved the SNR. The L-CAR operation was performed using GPU and added about 10% to the total processing time (Figure 7B).

After the slope filtering and L-CAR operations, the conditioned signals are stored in a disk or cached in RAM to expedite the successive data I/O operations. Although the conditioned signals are stored in a time-differentiated format, smoothed and non-differentiated waveforms can be quickly obtained by integrating the slope waveforms.

### Spike event detection and feature extraction

An ideal spike sorting algorithm should correctly detect spikes without false negatives or positives, and precise timing should be assigned to each spiking event. For HD probes, a spiking event is generally detected by several adjacent sites, and thus coincident spikes across multiple sites need to be grouped together to prevent multiple counting. For each spiking event, we extract the waveform information within a fixed-size spatiotemporal window (1 ms and 50 μm radius) centered at the largest spike-containing site. We find that 50 μm radius (10 nearest sites for the Neuropixels probes) provide a good balance between including most spikes from the same unit while excluding spikes from other units. The fixed-size window allows us to standardize a set features that we used to compare two spiking events regardless of their amplitudes or the spatial extent. Further, the fixed spatial template includes the sites containing sub-threshold spikes, which still carry useful information but have been previously discarded by a mask-based event definition used by others.

Initially, spike detection is carried out independently by each channel, and the spike timing is found by searching for the negative turning points below the detection threshold. The detection threshold is defined relative to the background noise activity (6x MAD) [7]. Next, each spike is compared to all other spikes found within its spatiotemporal neighborhood (±1 ms refractory period; <50 μm radius), and we only preserve the spikes having the absolute maximum amplitude within its local neighborhood. The remaining spikes are now referred as the event centers, and the waveforms surrounding the event centers are extracted to compute the set of low-dimensional features for clustering. For the event centers located near the either ends of the probe, we extract the waveforms from *K* number sites from the top or bottom (*K*=10 for the Neuropixels probe).

Principal component analysis (PCA) is widely used to extract low-dimensional features in conventional spike-sorting software, since it captures the waveform information unique to different units. However, PCA scores are sensitive to the noise-induced time shifts between the waveform templates and the spike waveforms. We circumvent this “time-jitter” problem by generating waveform templates unique to each spiking event. A template waveform is generated for each spiking event by normalizing the center site’s waveform, which has the highest SNR and has a fixed timing relationship amongst its neighboring sites’ waveforms. We accurately estimate the spike amplitudes by computing the covariance between the self template and the waveforms from each site. The self template is projected to the waveforms from each sites after the mean subtraction. The projection (or dot-product) operation cancels uncorrelated noise between sites, thus providing a more accurate estimation of the spike amplitudes than other measures such as peak-to-peak or peak amplitudes. We now refer to our projection-based feature as the covariance feature.

It is possible to extract more than one feature per site to increase the number of features available for clustering. For a given spiking event, we commonly observe a time-shifted version of spike waveforms in adjacent sites that are delayed by a few samples (~0.1 ms) relative to the center site containing the largest waveform. Since the temporal relationship between neighboring sites remain constant for a given unit, we capture this information by projecting a time-delayed (0.1 ms) version of the self template. The spike sorting accuracy improved after adding this secondary feature with a time delay.

### Position tracking and drift correction

We estimate the position of spiking event for the purpose of spike sorting and probe drift correction. Physical location of a spiking event provides valuable information about the identity of a neuron since relative positions between neurons remain mostly constant during a recording session (often lasting hours). Even so, it is common to observe physical movement between the probe and brain, which causes a gradual and systematic change in the spike amplitudes over time as neurons move closer or away from recording sites. Such probe drift can cause major errors in spike sorting if left uncorrected. We address this issue by estimating the location of each spike while detecting and correcting for the probe-wide movement.

The centroid position of a spiking event is calculated from the average of the site positions, and each site is weighted by its squared spike amplitude [15]. The spiking position is used to restrict the search range of its neighbors during clustering. This allows an efficient and accurate clustering since spikes from the same neuron form a tight spatial distribution in the absence of probe drift. Once the spike positions are determined, we check for a global drift in spike positions by comparing spatial relationships between adjacent time bins. We first generate a spatial representation of the spiking activity (activity map) by plotting the histogram of spike count for each spatiotemporal bin (2 um, 0.5 s bin size). We only include spikes with amplitudes above the median threshold, since small-amplitude spikes produce less accurate position estimation. An activity map of this kind reveals multiple bands of spiking activities at various depths and it becomes easier to spot global drifts. We automatically track probe drift over time by computing the cross-correlations between adjacent time bins (+/- 2 bins). The cross-correlation provides an optimal spatial shift between two time slices to match the spiking activity profile. We first run the cross-correlation using a smaller bin size (0.5 s) to determine and correct for the shift and then repeat the runs by doubling the time bin size until the bin size reaches the recording duration.

### Automated clustering

Clustering operations group similar spikes together to determine the unique identity of each neuron that generates a given spike. This is done in conventional spike sorting software [Klustakwik] by fitting a Gaussian mixture model (GMM) in the feature space. However, an observed distribution can deviate from the Gaussian assumption when a drift is present or if slow variations in the spike waveform occur. These non-ideal conditions lead to elongation of the distribution which results in false cluster splits. Further, the fitting procedure does not scale well with the increasing number of clusters and data generated by HD probes. The algorithmic complexity of the GMM fitting procedure, based on the expectation maximization (EM), is O(*d*×*l* + *k*×*l*^2^), where *d* is the number of dimensions, *l* is number of spikes and *k* is number of clusters. This means processing time for an ever increasing amount of data is untenable using this or similar algorithms.

We overcome these issues by applying a newly developed density-based clustering algorithm titled “Clustering by fast search and find of density peaks” (DPCLUS) [17]. Density-based clustering algorithms such DPCLUS uses pairwise distances between neighboring points to determine the cluster membership, and do not require the data to follow any particular distribution. The algorithm computes a local density (*ρ*) around each point representing a spiking event and the distance (*δ*) to the nearest point having a greater density, which emulates a gradient ascent operation. The local density is determined by the number of neighboring points within a cutoff distance (*d*_*c*_), which is automatically determined prior to the clustering operation. The dc parameter is set equal to 2% quantile of the distance distribution from a uniformly subsampled population of spikes. Once the rho and delta parameters are calculated, we linearly detrend the *δ vs. ρ* plot in the log-log scale and apply a threshold to find outliers in *δ* dimension. These outliers correspond to the local density peaks that are sufficiently far from other points at the same density level. These density peaks become cluster centers and cluster membership is copied to the nearest point with a lower density (gradient descent). This membership assignment repeats in the decreasing order of density until all points are assigned to clusters.

Density-based clustering algorithms require a pairwise distance calculation that scales quadratically to the number of spikes *O(n*^*2*^). We modified this algorithm to scale quasi-linearly to the number of spikes *O(n)* by taking an advantage of the fact that spikes from the same neuron come from close physical locations. We restrict the pairwise distance calculation to spikes that are located within half a vertical site spacing (10 μm), such that spiking events that are too far apart to originate from the same neuron will be excluded from the density (*ρ*) and distance (*δ*) calculations. Once we obtain the clustering output, we merge cluster pairs if they have similar average waveforms exceeding a correlation score of 0.975 [19]. At the end of clustering, we discard clusters having spikes less than the number of dimensions (*n*=20; two dimensions/site × 10 sites) and the clusters are sorted according to their center-of-mass locations [15].

### Manual verification

Despite the field’s best efforts to build an automated pipeline, algorithms can still make rare mistakes which require human operators to verify the machine-generated output. To accommodate this, we have built an interactive user interface (UI) that outputs a visualization of various information about individual units (putative neurons), and the relationship between multiple units. It is important to provide a responsive and intuitive UI such as this to assist users in performing efficient browsing, merging and splitting operations. Our UI offers semi-automated guides to save users’ time by suggesting the most optimal merging or splitting opportunities. It is important to enable the easy navigation and efficient browsing of very large datasets, especially for the HD probes that typically yield hundreds of units. We achieved a scalable sub-second user response time for a very large dataset (10s GB) by precomputing and caching frequently accessed information. In addition to visualizing the neural data, our interface serves as a common platform to integrate the behavioral data by visualizing the neural responses to various sensory input and motor output.

In terms of user interactions, users can browse the units sorted by their center-of-mass locations along the probe length by zooming and panning with a mouse. Users can select a unit of interest with a left mouse click to display its mean and individual spike waveforms, physical distribution of the signal amplitude, inter-spike intervals (ISI) and cross-correlations with other nearby units. Users can also compare the relationship between two units by selecting the second unit with a right mouse click to superimpose their waveforms, ISI, return maps and spike center-of-mass positions. Our UI additionally offers a time vs. amplitude view to display information about any detected probe drift. Users can merge two units if their features are continuous in time, or split in the time view if two units are incorrectly joined by crossing over each other. It is possible to quickly inspect neural tuning properties by generating peristimulus time histograms (PSTH), spike rasters and place field maps from given behavioral data. Our UI provides keyboard shortcuts for frequently used operations such as merging, splitting, or deleting clusters. Additional operations from those listed above are available from the menu bar.

### Hardware and software platform

We built our spike sorting pipeline for a single workstation environment with a high-end GPU (graphical processing unit). We’ve taken advantage of a local parallel computing environment that utilizes multiple CPU and GPU cores, while the data input/output (I/O) operations are restricted to the same local machine for faster access. This computing paradigm has several advantages over a cluster computing approach which, while popular for some big data applications, is not ideal for spike sorting operations. Cluster or cloud computing platforms add considerable speed bottlenecks by transmitting a large volume of data over a network, which is an order of magnitude slower than a local data I/O speed. Spike sorting performance on a large (~10s GB) dataset is largely determined by efficient data management and parallel computations.

We found significant speed enhancements by installing sufficient RAM (128 GB) to cache the data and high-speed data storage (SSD or RAID). While solid-state drives (SSD) can offer up to four times the speed of a single hard drive, they offer much lower storage volume per cost. It is important to consider a long-term strategy for managing tens of TB of neural recordings from HD probes, since it is expected to accumulate hundreds of gigabytes of recordings per day from several animals. We have successfully used locally-attached hard drive arrays that provide a fast read/write speed (~1 GB/s with Thunderbolt II interface) while providing sufficient storage volume (up to 80 TB; 10 TB/disk × 8 disks). We opted to use RAID-5 configuration to protect against a single hard drive failure at a cost of a single drive storage capacity.

Also contributing to our fast and scalable performance was our writing optimized algorithms that take advantage of the aforementioned massively parallel local computing paradigm. We made an extensive use of GPU for computation-intensive tasks and multiple CPU cores for data-driven tasks. GPUs can offer overall speed gains over CPUs if the time savings from GPU computation is greater than the time delays from transmitting data between the GPU memory (GRAM) and system memory. To optimize data transmission in this way we reduced unnecessary I/O operations by caching repeatedly accessed information in RAM or GRAM (GPU memory). Several high-end GPUs offer enough memory (12 GB for NVIDIA Titan X) to store the entire recording duration from several neighboring sites. GPUs also offer a much greater number of cores (~3000) than CPUs (~10), and GPU threads from the same core (=stream processor) can collectively access a small (~48 KB) yet fast on-chip cache (shared memory). We wrote our clustering code in a low-level, hardware-specific C-language (CUDA) to take a full control of the GPU hardware architecture. This GPU-centric approach led to our clustering engine achieving a significant speed gain (70x) over CPU-based code (Figure 7C).

While low-level languages used in the way described above allow greater control of hardware and enable faster execution speed, they generally take much greater effort to develop, test and improve the code. We therefore built the rest of our spike sorting pipeline on a high-level language platform (MATLAB) to take advantage of many built-in supports for scientific computations. Our pipeline uses signal processing and parallel computing MATLAB toolboxes, which offer seamless integration with local parallel computing hardware as well as a simple interface between low level and high level languages. Although such a platform is not free, the computational libraries are maintained, documented and optimized by professional developers, and it has a broad user base in the neuroscience community.

### Sorting performance measurements

The goal of our sorting platform is to devise a general algorithm that works accurately across many datasets. The quality of our sorting is to be evaluated without changing the sorting parameters since we do not want to overfit our parameters to match a particular ground truth dataset. In addition to using the experimentally obtained ground-truth datasets, we also generated simulated ground truth datasets which closely emulates other ground truth datasets obtained from real recordings. We simulated several types of probe geometries with site spacing (20-32 μm center-to-center) comparable to the real ground truth datasets. Our choice of algorithms and parameters were guided by the overall performance across all datasets tested. We used a common set of parameters across all datasets except for the recording-specific parameters such as the probe layout and the recording file format.

### Ground truth datasets

It is important to measure the spike sorting performance with known neuronal identities associated with each spike, or ground truth information as commonly referred. It would be ideal to measure the intracellular voltages from all biological neurons within the detection volume, but such ground truth information is yet to exist. Therefore, we combine three types of currently available ground truth datasets, each with strength and weakness, to validate our spike sorting performance in a comprehensive manner. It is possible to generate a ground-truth information for a single neuron by simultaneous recordings from a micropipette and extracellular electrodes. Intra- or juxtra-cellular recordings provide precise spike timing information to be compared with the spike sorting results from extracellular electrodes. However, the paired recording technique is currently limited to one neuron at a time, and it is technically challenging to obtain a paired recording within the same detection volume (<75 μm) due to difficulties with precisely positioning a micropipette near a silicon probe.

### Hybrid ground truth

Alternatively, it is possible to generate ground truth datasets with higher throughput by manually isolating a subset of well-separated units and inserting their spikes at predetermined time and probe locations. This approach is commonly referred to as a hybrid ground truth [9,19]. Spike waveforms are denoised by applying singular value decomposition (SVD) prior to adding the ground truth spikes, in order to preserve the SNR afterwards. This type of ground truth is referred as the *hybrid ground truth* since it requires a hybrid of human judgement and automated algorithm to generate such dataset. Although the hybrid ground truth offers more number of identified units per recording session than the ground truth from paired recordings, only a small subset of well-isolatable units can be tested. This method is biased toward testing high-amplitude units that can be sorted reliably, since lower amplitude units near the threshold cannot be manually sorted reliably.

### Simulated ground truth

In order to quantify the sorting performance across wide range of unit amplitudes, we generated ground truth by simulating a population of biophysically realistic neurons. It is now possible to simulate a population of neurons in a cortical column that faithfully captures individual ion-channel dynamics. Our biophysical simulation randomly distributed ~710 neurons within a sensing volume of the probe (200×200×600 μm), and each neuron is constructed from real morphology of various cell types. Our simulation closely matches the real recordings in terms of the spatial spread of the spike waveforms (Figure 5B,D), and the identity of all units and their spike timing information are precisely determined from their simulated intracellular voltage traces.

Biophysically realistic network simulations were setup in order to generate simulated ground truth extracellular depth recordings datasets. The main engine used for these biophysical simulations was NEURON version 7.4 [31] used with custom-written python (version 2.7) wrapper-algorithms that were used to define, setup and instantiate network models and simulations as well as save output.

The network model consists of two cell types, excitatory pyramidal neurons (2,560) and inhibitory basket cells (640). Pyramidal single-neuron models used in our network were published by Hay and co-workers as part of their study on models capturing a wide range of dendritic and perisomatic active properties [32]. These single-neuron models were shown to capture a number of intracellular characteristics such as backpropagating action potentials, dendritic Calcium electrogenesis, etc. In addition, a later study showed that these computational models accurately capture the extracellular signature of neocortical pyramidal neurons [18]. We adopted inhibitory basket cell models from Hu et al. [33] who developed these models to capture a number of features such as backpropagation and dendritic Na/K ratio. Notably, pyramidal and basket cell models had active dendrites, a feature critical for capturing extracellular spiking activity [1,13,18,34]. Each neuron received external excitatory synaptic input emulating AMPA synapses. The number and timing of these external synaptic inputs was set such as to reach specific spike frequency output.

With regards to calculating the simulated extracellular recordings, the so-called line source approximation [1] was used to estimate the extracellular voltage at various locations assuming a perfectly homogeneous, isotropic and purely resistive (0.3 S/m) extracellular space [18,35–38]. Specifically, the Neuropixels probe electrode location was adopted from layout specifications. Network simulations were performed at the San Diego supercomputer facility via the NSG portal.

### Ground truth from paired recordings

We used two separate paired-recording experiments (*in-vivo* and *in-vitro*) to collect four cell pairs that exceeded our selection criteria (8x SNR or 2x detection threshold). First, the *in-vitro* paired recording experiment from a brain slice is described in detail in [18]. Briefly, whole-cell patch recordings of excitatory and inhibitory neurons in rat somatosensory cortex slice were performed while positioning a silicon probe (8 sites/shank, H32, Neuronexus) in their vicinity to concurrently record intra- and extracellular voltages. Individual neurons were stimulated via intracellular DC current injections and intracellular somatic membrane potential was recorded during spiking simultaneously with the extracellular potential at various locations along the silicon probe. Due to the lack of spontaneous spiking activities except for the one that was injected with current, we combined ten separate recording sessions and sorted them together. We used two ground-truth units that exceed our selection criteria.

We also included *in-vivo* juxtacellular recordings generated by Neto et al. [10]. They used a similar methodology to generate simultaneous juxta- and extracellular recordings in the motor cortex of an anesthetized rat. The extracellular recordings were obtained using silicon probes containing 32 or 127 sites per shank with 22.5-25 μm site spacing. We used two recording sessions that exceeded our selection criteria.

## Acknowledgements

CM, SLG and CAA would like to thank NSG portal personnel for offering access to the San Diego supercomputing facility as well as for troubleshooting and support. SLG and CAA thank the Allen Institute founders, P. G. Allen and J. Allen, for support. The authors thank Drs. A. Lee, K. Svoboda and K. Harris for helpful discussions, and Drs. N. Steinmetz, J. Neto and A. Kampff for sharing the ground-truth datasets. JJJ, CL and TDH received support from the Howard Hughes Medical Institute.

## References

1 Gold C, Henze DA, Koch C, Buzsáki G. On the origin of the extracellular action potential waveform: A modeling study. J Neurophysiol. 2006;95: 3113–3128.

2. Lopez CM, Andrei A, Mitra S, Welkenhuysen M, Eberle W, Bartic C, et al. An Implantable 455-Active-Electrode 52-Channel CMOS Neural Probe. IEEE J Solid-State Circuits. 2014;49: 248–261.

3. Scholvin J, Kinney JP, Bernstein JG, Moore-Kochlacs C, Kopell N, Fonstad CG, et al. Close-Packed Silicon Microelectrodes for Scalable Spatially Oversampled Neural Recording. IEEE Transactions on Biomedical Engineering. 2016;63: 120–130.

4. Lopez CM, Mitra S, Putzeys J, Raducanu B, Ballini M, Andrei A, et al. 22.7 A 966-electrode neural probe with 384 configurable channels in 0.13µm SOI CMOS. 2016 IEEE International Solid-State Circuits Conference (ISSCC). 2016. doi:10.1109/isscc.2016.7418072

5. Harris, T.D., Jun, J.J., Karsh, B., Barbarits, B., Sun, W.L., Musa, S., Putzeys, J., Andrei, A. & Lopez, C. The neuropixels project: design and performance of 966 selectable site 384 channel Si probes. 2016 Neuroscience Meeting Planner San Diego, CA. Society for Neuroscience; p. Program No. 774.07. Online.

6. Seymour JP, Wu F, Wise KD, Yoon E. State-of-the-art MEMS and microsystem tools for brain research. Microsystems & Nanoengineering. Nature Publishing Group; 2017;3: 16066.

7. Quiroga RQ, Quian Quiroga R, Nadasdy Z, Ben-Shaul Y. Unsupervised Spike Detection and Sorting with Wavelets and Superparamagnetic Clustering. Neural Comput. 2004;16: 1661–1687.

8. Rossant C, Kadir SN, Goodman DFM, Schulman J, Hunter MLD, Saleem AB, et al. Spike sorting for large, dense electrode arrays. Nat Neurosci. 2016;19: 634–641.

9. Pachitariu M, Steinmetz N, Kadir S, Carandini M, Harris KD. Kilosort: realtime spike-sorting for extracellular electrophysiology with hundreds of channels [Internet]. bioRxiv. 2016. p. 061481–1. doi:10.1101/061481

10. Neto JP, Lopes G, Frazão J, Nogueira J, Lacerda P, Baião P, et al. Validating silicon polytrodes with paired juxtacellular recordings: method and dataset. bioRxiv. 2016; doi:10.1101/037937

11. Hawrylycz M, Anastassiou C, Arkhipov A, Berg J, Buice M, Cain N, et al. Inferring cortical function in the mouse visual system through large-scale systems neuroscience. Proc Natl Acad Sci U S A. 2016;113:7337–7344.

12. Ludwig KA, Miriani RM, Langhals NB, Joseph MD, Anderson DJ, Kipke DR. Using a common average reference to improve cortical neuron recordings from microelectrode arrays. J Neurophysiol. 2009;101:1679–1689.

13. Schomburg EW, Anastassiou CA, Buzsáki G, Koch C. The spiking component of oscillatory extracellular potentials in the rat hippocampus. J Neurosci. 2012;32: 11798–11811.

14. Kadir SN, Goodman DFM, Harris KD., High-dimensional cluster analysis with the masked EM algorithm. Neural Comput. 2014;26: 2379–2394.

15. Nádasdy Z, Csicsvari JO, Penttonen MA, Hetke JA, Wise KE, Buzsaki G. Extracellular recording and analysis of neuronal activity: from single cells to ensembles. In: Eichenbaum HB, Davis JL, editors. Neuronal Ensembles: Strategies for Recording and Decoding. Wiley-Liss; 1998. pp. 17–55.

16. Harris KD, Henze DA, Csicsvari J, Hirase H, Buzsáki G. Accuracy of tetrode spike separation as determined by simultaneous intracellular and extracellular measurements. J Neurophysiol. 2000;84: 401–414.

17. Rodriguez A, Laio A. Clustering by fast search and find of density peaks. Science. 2014;344: 1492–1496.

18. Anastassiou CA, Perin R, Buzsáki G, Markram H, Koch C. Cell type- and activity-dependent extracellular correlates of intracellular spiking. J Neurophysiol. 2015;114: 608–623.

19. Yger P, Spampinato GLB, Esposito E, Lefebvre B, Deny S, Gardella C, et al. Fast and accurate spike sorting in vitro and in vivo for up to thousands of electrodes [Internet]. 2016. doi:10.1101/067843

20. Wu F, Stark E, Im M, Cho I-J, Yoon E-S, Buzsáki G, et al. An implantable neural probe with monolithically integrated dielectric waveguide and recording electrodes for optogenetics applications. J Neural Eng. 2013;10: 056012.

21. Wu F, Stark E, Ku P-C, Wise KD, Buzsáki G, Yoon E. Monolithically Integrated μLEDs on Silicon Neural Probes for High-Resolution Optogenetic Studies in Behaving Animals. Neuron. 2015;88: 1136–1148.

22. Hoffman L, Subramanian A, Helin P, Bois BD, Baets R, Dorpe PV, et al. Low Loss CMOS-Compatible PECVD Silicon Nitride Waveguides and Grating Couplers for Blue Light Optogenetic Applications. IEEE Photonics J. 2016;8: 1–11.

23. Welkenhuysen M, Hoffman L, Luo Z, De Proft A, Van den Haute C, Baekelandt V, et al. An integrated multi-electrode-optrode array for in vitro optogenetics. Sci Rep. 2016;6: 20353.

24. Shim E, Chen Y, Masmanidis S, Li M. Multisite silicon neural probes with integrated silicon nitride waveguides and gratings for optogenetic applications. Sci Rep. 2016;6: 22693.

25. Zhang F, Gradinaru V, Adamantidis AR, Durand R, Airan RD, de Lecea L, et al. Optogenetic interrogation of neural circuits: technology for probing mammalian brain structures. Nat Protoc. 2010;5: 439–456.

26. Zhao S, Ting JT, Atallah HE, Qiu L, Tan J, Gloss B, et al. Cell type-specific channelrhodopsin-2 transgenic mice for optogenetic dissection of neural circuitry function. Nat Methods. Nature Research; 2011;8: 745–752.

27. Ledochowitsch P, Yazdan-Shahmorad A, Bouchard KE, Diaz-Botia C, Hanson TL, He J-W, et al. Strategies for optical control and simultaneous electrical readout of extended cortical circuits. J Neurosci Methods. 2015;256: 220–231.

28. Ledochowitsch, P., Ducros, M., Liu, R., Buice, M.A., Mitelut, C., Anastassiou, C., Saggau, P., Blanche, T.J.}. Linking electrophysiology and optophysiology in vivo. Neuroscience Meeting Planner, Chicago, IL: Society for Neuroscience, 2015. 2015. p. Program No. 598.26. Online.

29. Allen, B., Moore-Kochlacs, C., Scholvin, J., Kinney, J., Bernstein, J., Kodandaramaiah, S.B., Kopell, N., Boyden, E. Towards ground truth in ultra-dense neural recording. 2016 Neuroscience Meeting Planner San Diego, CA. Society for Neuroscience; p. Program No. 774.09. Online.

30. Voigts, J., Cuevas-Lopez, A., Newman, J.P., Siegle, J.H., Lopes, G.C., Mondragon, S.L., Patel, Y.A., Burguière, E., Kemere, C., Moore, C.I., Widge, A.S., Kampff, A., ViventiI, J., Wilson, M. A open-source system and interface standards for very high data rate neurophysiology and low latency closed-loop experiments. Neuroscience Meeting Planner San Diego, CA: Society for Neuroscience, 2016. 2016. p. Program No. 848.13. Online.

31. Hines ML, Carnevale NT. The NEURON Simulation Environment. Neural Comput. 1997;9: 1179–1209.

32. Hay E, Hill S, Schürmann F, Markram H, Segev I. Models of neocortical layer 5b pyramidal cells capturing a wide range of dendritic and perisomatic active properties. PLoS Comput Biol. 2011;7: e1002107.

33. Hu H, Martina M, Jonas P. Dendritic mechanisms underlying rapid synaptic activation of fast-spiking hippocampal interneurons. Science. 2010;327: 52–58.

34. Reimann MW, Anastassiou CA, Perin R, Hill SL, Markram H, Koch C. A biophysically detailed model of neocortical local field potentials predicts the critical role of active membrane currents. Neuron. 2013;79: 375–390.

35. Logothetis NK, Kayser C, Oeltermann A. In vivo measurement of cortical impedance spectrum in monkeys: implications for signal propagation. Neuron. 2007;55: 809–823.

36. Anastassiou CA, Perin R, Markram H, Koch C. Ephaptic coupling of cortical neurons. Nat Neurosci. 2011;14: 217–223.

37. Anastassiou CA, Buzsaki C, Koch C, Quiroga R, Panzeri S. Biophysics of extracellular spikes. Principles of Neural Coding. CRC Press Boca Raton, FL; 2013;15: 146.

38. Anastassiou CA, Shai AS. Psyche, Signals and Systems. In: Buzsáki G, Christen Y, editors. Micro-, Meso- and Macro-Dynamics of the Brain. Springer International Publishing; 2016. pp. 107–156.

